# Immunogenicity and efficacy of Heat Inactivated Whole-Cell Vaccine to increase murine survival after Extensively drug-resistant *Acinetobacter baumannii* infection

**DOI:** 10.1101/2021.08.30.458307

**Authors:** Nazmun Sharmin, Mahbub E Khoda, SM Shamsuzzaman, Mohammad Nazim Uddin

**Affiliations:** Department of Microbiology, Dhaka Medical College, Dhaka, Bangladesh; Department of Pulmonology, Dhaka Medical College Hospital, Dhaka, Bangladesh; Ministry of Health, Saudi Arabia

**Keywords:** Heat Inactivated Whole Cell Vaccine, *A. Baumannii*, XDR, Serum Bactericidal Antibody

## Abstract

Due to the rapid emergence of extensively drug-resistant (XDR) strains worldwide there is a necessity for greater consideration for the role of preventive vaccines to combat these pathogens. The study reported the effectiveness of a heat-inactivated whole-cell vaccine (HI-WCV) against *A. baumannii* to produce protective immunity with evaluation of the bactericidal antibody responses in mice after intramuscular inoculation with different XDR strains of *A. baumannii*. Six XDR *A. baumannii* strains emulsified with complete freund’s adjuvant (CFA) were used for intramuscular inoculation in the experimental group of mice (n=4) in three different phases on 14 days interval whereas placebo-controlled mice (n=4) received phosphate-buffered saline emulsified with CFA in same the schedule. Serum was collected from tail blood of each mouse on 10^th^ day after each inoculation and by cardiac puncture on 14^th^ day after lethal dose from the experimental group and after death of the placebo-controlled group. Mice inoculated with heat-inactivated whole-cell vaccine survived and showed a higher neutralizing (IgG) antibody response after 2nd (>7-fold titre) and 3rd inoculation (>12-fold titre) whereas all mice from placebo-controlled group did not survived. Sera from experimental group of mice collected (38^th^ day) after 3rd inoculation resulted in higher serum bactericidal killing index (42.12) after three hours incubation with an XDR *A. baumannii* over sera collected after 1st and 2nd inoculation (50% cutoff value=14.5). These results suggest that intramuscular vaccination with heat-inactivated whole-cell vaccine stimulated IgG antibody responses that can increase murine survival after XDR *A. baumannii* infection.

## Introduction

*Acinetobacter* species are non-fermentative, non-fastidious, catalase-positive, oxidase negative, non-motile and strictly aerobic gram-negative coccobacilli (1). Among these species *A. baumannii* causes serious infections that are associated with high morbidity and mortality rates (2) particularly, in immunocompromised patients, elderly, premature neonates and patients that had recently undergone surgery or experienced a major trauma or were previously admitted to contaminated ICUs (3) causing septicemia, endocarditis, meningitis, skin and soft tissue infection, wound infection, respiratory tract infections and urinary tract infections (4). The World Health Organization declared that *A. baumannii* is one of the most serious ESKAPE organisms (*Enterococcus faecium, Staphylococcus aureus, Klebsiella pneumoniae, A. baumannii, Pseudomonas aeruginosa*, and *Enterobacter* species) that effectively escape the effects of antibacterial drugs (5). Now, MDR and XDR *A. baumannii* have emerged worldwide as a critical problem and have been increasingly reported (6) as a result of widespread and indiscriminate use of antibiotics.

*A. baumannii* exhibits an intrinsic resistance to a range of antibiotics such as amoxicillin, narrow spectrum cephalosporins, ertapenem, trimethoprim and chloramphenicol (7). In addition*, A. baumannii* has managed to acquire resistance genes against several classes of antibiotics through the transfer of plasmids, transposons and integrons from other gram-negative bacteria (8). Currently, there are a small number of drug candidates like colistin, tigecycline, eravacycline (9), cefiderocol and other therapeutic options such as monoclonal antibody (10), phages (11) that may be used to treat XDR *A. baumannii* infections. Several researchers adopted the idea of solving the *A. baumannii* problem by vaccination (12). However, to date, there are no licensed vaccines against XDR *A. baumannii*.

Among the inactivated vaccine trials there are some single antigen candidates that have been tested includes, the outer membrane protein (OmpA), the K1 capsular polysaccharide, the biofilm-associated protein (Bap) and the membrane associated polysaccharide (poly-N-acetyl-β-1-6-glucosamine) (13). Heat-inactivated whole-cell vaccines are easy to prepare, inexpensive and does not require expensive denaturation of antigens, that may induce conformational changes of the epitopes. previous studies have shown that, emulsification with CFA and IM immunization in mice generate hyper-immune antibodies against its multi-antigenic components, mostly those belonging to the outer membrane (14), providing diverse protection against several *A. baumannii* strains with a complete bacterial clearance from the infected tissues resulting in increased mice survival rates (15). Furthermore, the vaccine proved to induce a protective level of immunoglobulins and reduced the pro-inflammatory cytokines serum levels of IL-1β, TNF-α, and IL-6 that are normally associated with sepsis and helped to secure high survival rates in vaccinated mice associated with both pneumonia and sepsis (16) that could be life-saving in case of outbreaks or critical conditions (17).

We, therefore attempted to develop a HI-WCV derived from multiple strains of XDR *A. baumannii* after detecting several drug resistance genes and investigated potential efficacy of enhancing protective immunity against XDR *A. baumannii*, by evaluating the ability of the vaccine to improve the complement-mediated serum bactericidal activities of the immunized mice sera after intramuscular inoculation.

## Results

### *In vivo* assay

Immunized mice sera were collected before 1^st^ inoculation and on 10^th^ day after each vaccine inoculation from both experimental and placebo-controlled mice and humoral immune response was evaluated by the mean Optical Density (OD) values of duplicates of serum IgG absorbance at 450 nm in 1:160 dilution by indirect ELISA (**Figure 1B**). Among EG mice mean OD values of pre-inoculation and after 1^st^ inoculation (Day 10) was 0.29±0.028 and 0.330±0.002 respectively. After 2^nd^ inoculation (Day 24) mean OD value was 2.251±0.262 showing rise of serum antibody titre resulted in a rapid memory response peak (*P*<0.001). Moreover, the mean OD value was highest (3.487±0.275) after 3^rd^ inoculation where serum was collected on day 38 (*P*<0.001). However, there was a decrease in mean OD value (2.981±0.119) after lethal dose (Day 56) (*P*< 0.001). The sera from placebo-controlled mice contained very low levels of detectable antibodies at any time-point with an average OD value of 0.0376±0.012 which was very similar to NC mice serum OD value and all the mice from this group died within 24 hours after injection the lethal dose.

**Figure 1:**
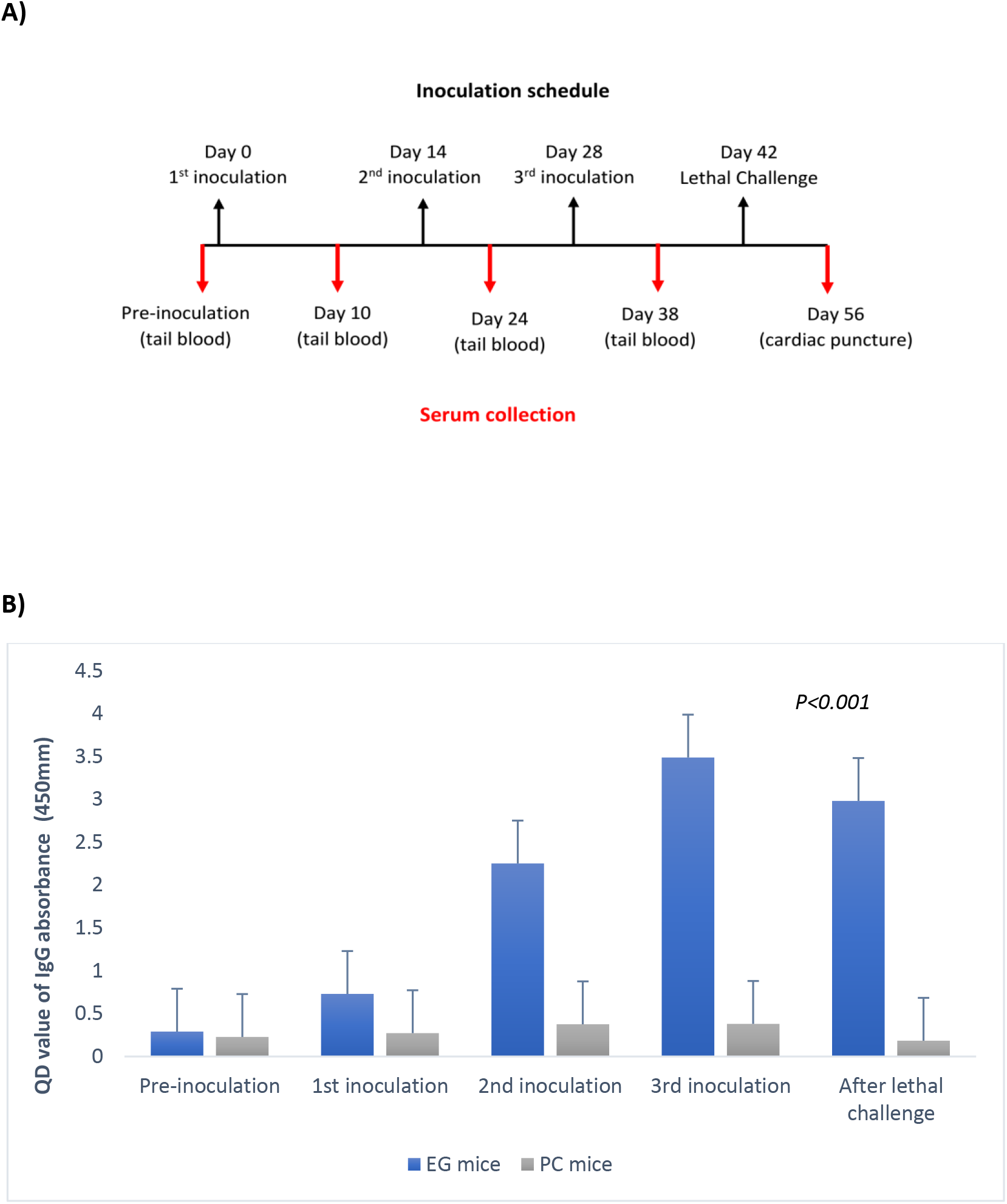
**A)** Both EG and PC group (n=4) was inoculated with HI-WCV on Day 0, 14, 28 intramuscularly and lethal challenge was given intraperitoneally on Day 42. Serum was collected from tail blood before inoculation and on Day 10, 24, 38. Blood was collected by cardiac puncture from euthanized NC group, immediately after death from PC group and on Day 56 after the lethal challenge from EG group. **B)** Mean Optical Density of serum samples before and after each inoculation (Day 0, 14, 28) and after lethal challenge (Day 42) of both EC Group and PC Group of mice (n=4). Here, *P< 0.001* except in pre-inoculation (*P=2.77*).

### *In vitro* assay

Immunized mice sera collected on days 10, 24, 38 and 56 were tested for the ability to promote *in vitro* complement-mediated serum bactericidal activity of the test XDR *A. baumannii*. **Table 1** showed the organized colony counts per spot from Mueller Hinton agar plates after overnight incubation at 37°C at 8 different dilutions. Bacterial killing is clearly seen for all samples tested in the first three dilutions and as samples are diluted further up the plate, a decrease in bacterial killing is seen where serum is less concentrated. For control A the mean CFU count was 36 in all dilutions. For control B the mean CFU count was 29 in all dilutions. For pre-inoculation the mean CFU count was between 24 to 35 in all dilutions indicating no inhibition of bacterial survival. The calculated non-specific killing (NSK) was 23%. Here, 50% killing index (KI) cut off value was 14.5. The 1st, 2nd, 3rd inoculation and after lethal challenge the mean CFU count for 1st three dilution (1:8, 1:24, 1:72) at 25% and 50% guinea pig complement concentration was between 1.5 to 2.5, 0 to 2, 0 to 1.5 and 0 to 1.5 respectively, which indicated bactericidal efficacy of the immunized serums. The highest SBA KI 42.12 was found after 3^rd^ inoculation, whereas the lowest SBA KI 19.1 was found after 1^st^ inoculation. **Figure 2: a), b), c), d**).

**Figure 2:**
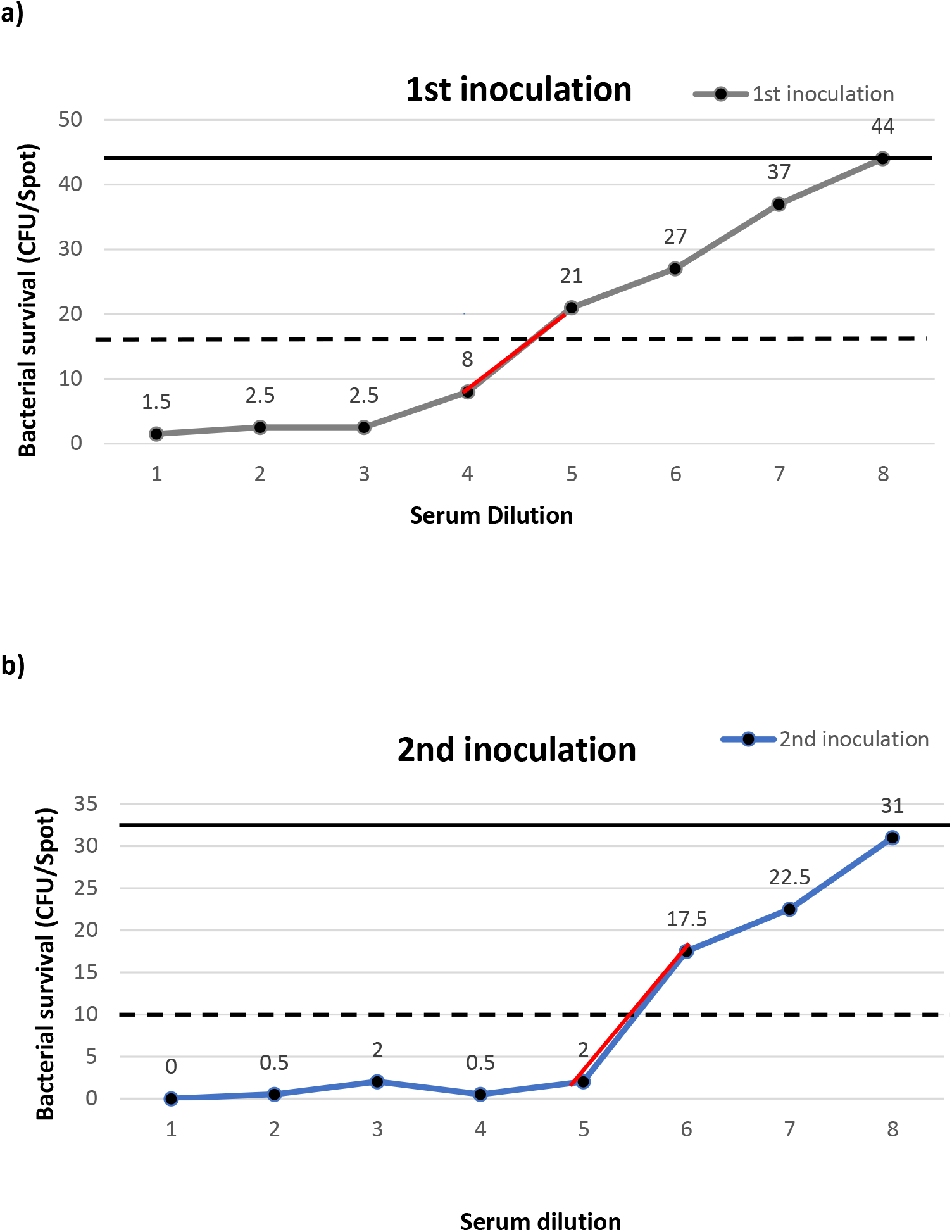
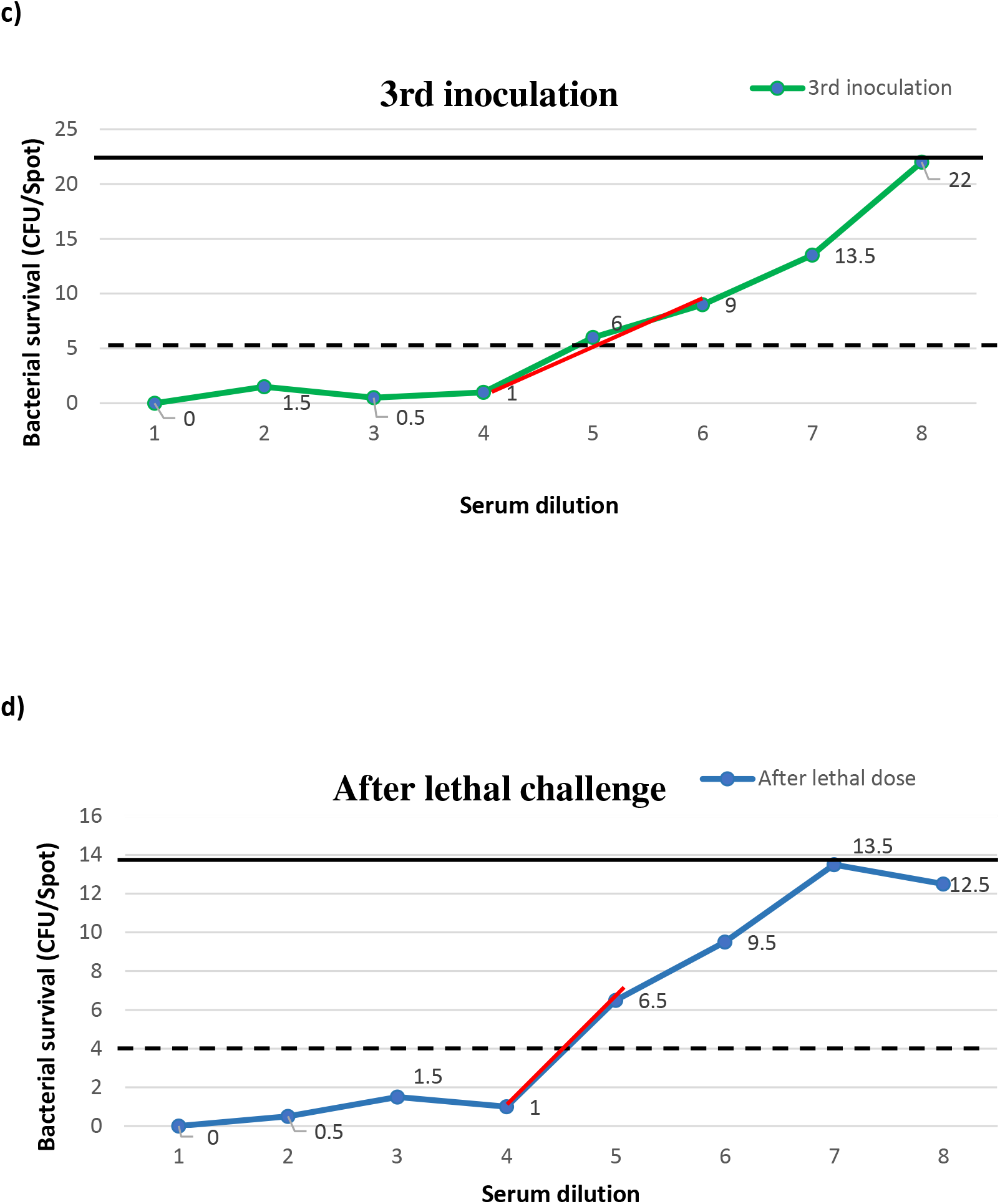
**a), b), c), d),** shows a schematic of linear interpolation after each inoculation. The number of surviving bacteria (y-axis) at each dilution of serum tested (x-axis) is plotted (balls) and individual points were connected by a thin line. The solid and dashed horizontal lines indicated 0% and 50% killing, respectively. The serum dilutions above (e.g., Dilution 5) and below (e.g., Dilution 4) the 50% killing line were connected by a red line and bactericidal titer (KI) was indicated.

**Table 1:**
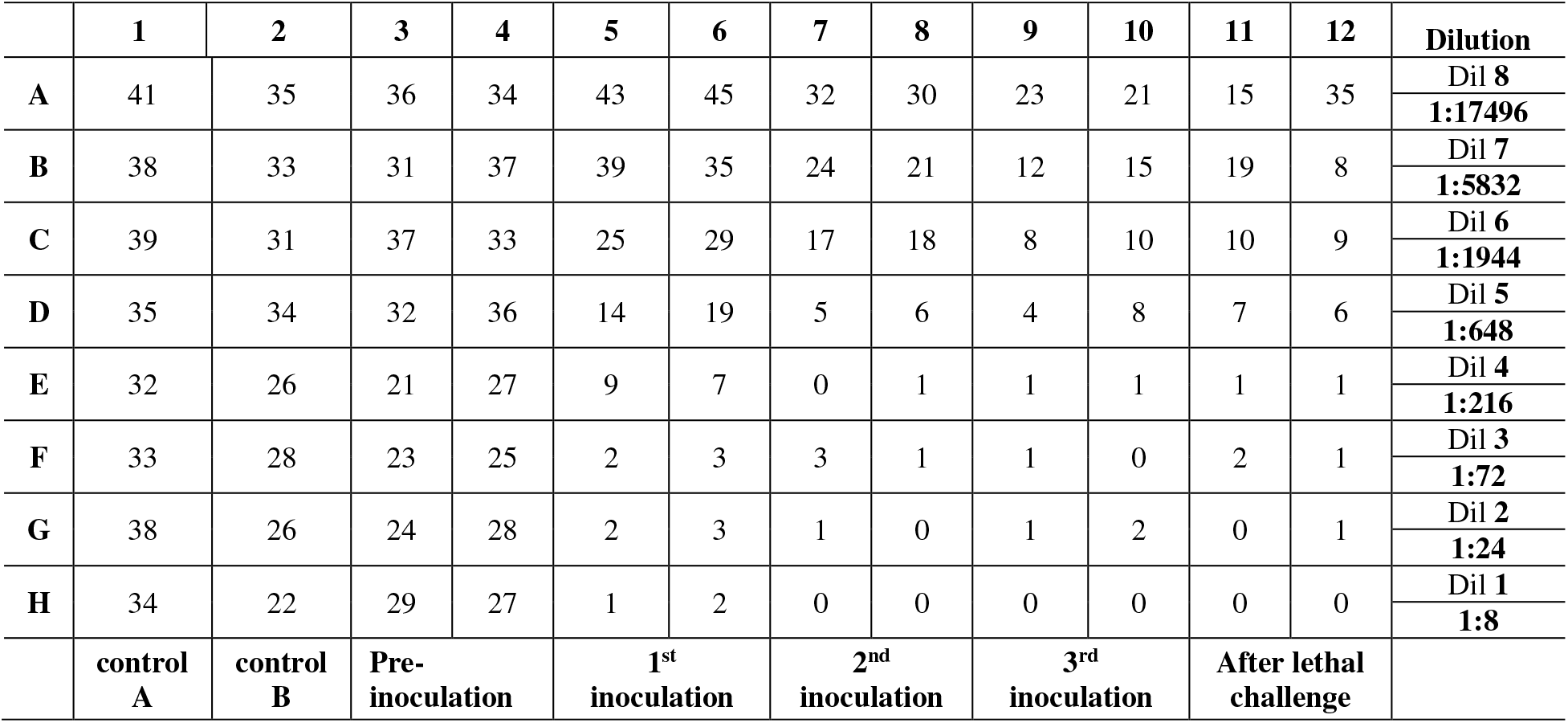
The organization of CFU counts/spot from Mueller Hinton agar plates

### Comparison of *in vivo* and in vitro assay

The neutralizing (IgG) antibody titer were significantly increased after 2nd (7-fold) and 3rd vaccination (12-fold). On the other hand, 7-fold rise in SBA KI from 50% SBA cutoff value was only shown after 3rd inoculation **(Table 2).**

**Table 2:**
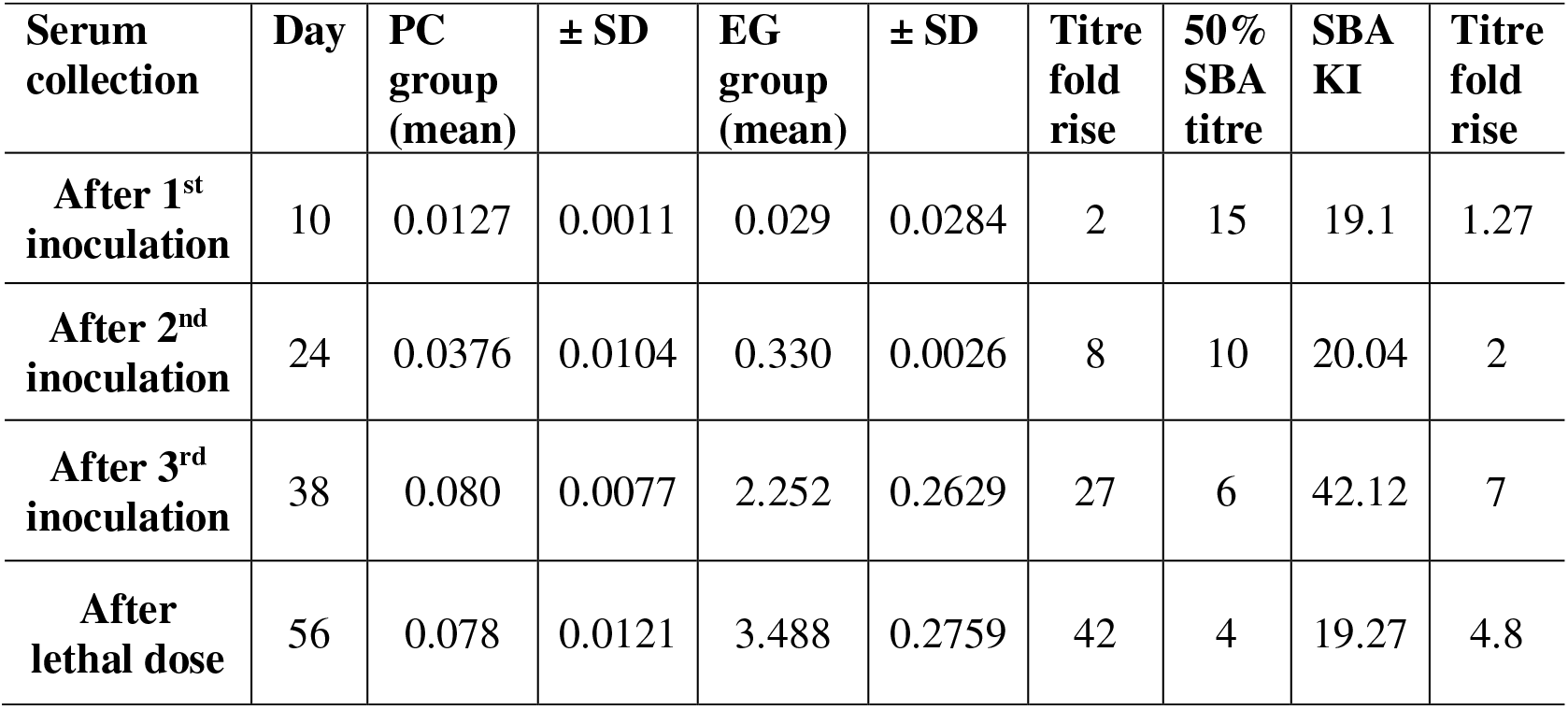
Comparison of the differences of rising in IgG antibody and SBA titers after vaccination between PC and EG group mice.

| **Serum collection** | **Day** | **PC group (mean)** | **± SD** | **EG group (mean)** | **± SD** | **Titre fold rise** | **50% SBA titre** | **SBA KI** | **Titre fold rise** |
| --- | --- | --- | --- | --- | --- | --- | --- | --- | --- |
| **After 1<sup>st</sup> inoculation** | 10 | 0.0127 | 0.0011 | 0.029 | 0.0284 | 2 | 15 | 19.1 | 1.27 |
| **After 2<sup>nd</sup> inoculation** | 24 | 0.0376 | 0.0104 | 0.330 | 0.0026 | 8 | 10 | 20.04 | 2 |
| **After 3<sup>rd</sup> inoculation** | 38 | 0.080 | 0.0077 | 2.252 | 0.2629 | 27 | 6 | 42.12 | 7 |
| **After lethal dose** | 56 | 0.078 | 0.0121 | 3.488 | 0.2759 | 42 | 4 | 19.27 | 4.8 |

## Discussion

Over the past few decades, *A. baumannii* has emerged as a major healthcare facility acquired bacterial infection because of resisting many classes of antibiotics by virtue of chromosome-mediated genetic elements (18), leading to rapid emergence of MDR, XDR and PDR strains that causes the consideration for preventive vaccines to combat. Though some single antigen candidates have been tested, these are still under trial. Various studies regarding heat-inactivated whole-cell vaccines showed that these were easy to prepare, inexpensive and did not require expensive denaturation of antigens, that may induce conformational changes of the epitopes and produced protective immunity (IgG Ab) against XDR *A. baumannii*, in murine models after intramuscular administration (19). In the current study, sera collected from each mouse after each inoculation was analyzed for IgG antibody absorbance by indirect ELISA at 450 nm. The OD values of serum IgG absorbance at 1:160 dilution showed that there was significant rise (*P*<0.001) in OD value of IgG antibody absorbance within the EG group of mice after 2nd (day 14) and 3rd (day 28) inoculation though the level was slightly decreased after intraperitoneal injection (day 42) of lethal dose with 1.5×10^8^ CFU/ml live XDR *A. baumannii* which might be due to utilization of the previously formed IgG antibody to neutralize *A. baumannii* antigen. The sera from PC mice contained very low levels of detectable antibodies after all inoculation with an average OD value of 0.0455±0.03244 which was very similar to NC group mice. Hence the mice from the EG group survived where all the mice from PC group were dead after lethal challenge as the serum neutralizing antibody was insufficient for them to survive.

In a study by Shu *et al*. (2016) in Malaysia showed that, heat inactivated whole-cell vaccine immunized mice sera collected on days 0 to 36 had an increase in IgG antibody. The OD values of IgG absorbance of immunized mice sera collected after second inoculation on day 22 resulted in a rapid memory response peak. However, the OD values on day 36 in immunized mice sera, showed a slight decrease in the antibody response. The sera from placebo-inoculated control mice contained very low levels of detectable antibodies at any time-point with an average OD value of 0.30±0.03 which was quite similar with the current study. The findings suggested that inoculation with heat inactivated whole-cell vaccine stimulated high levels of serum IgG antibody recognizing the XDR *A. baumannii* strain as the target antigen.

Activation of the complement pathway is an important immune defense mechanism present in serum. Therefore, complement activation could play an important role in minimizing the ability of *A. baumannii* to escape immune clearance. The present study, showed that the sera obtained from mice inoculated with heat-inactivated whole-cell vaccine were able to activate complement-mediated bacteriolysis in the presence of 25% guinea pig complement at 1:216 serum dilution after three hours of incubation with XDR *A. baumannii*. In this study, the calculated 50% SBA cutoff value was 14.5. It was noted that the bacteriolysis effects in sera of mice from the 1st and 2nd inoculation with HI-WCV collected on 10^th^ day (SBA KI=19.1) and 24^th^ day (SBA KI=20.04) showed only a slight increase from the 50% SBA titre, suggesting that sera from a two subsequent immunization did not significantly increase the complement-mediated serum bacteriolysis activity. However, the SBA KI (42.12) of the EG group (n=4) after 3rd inoculation (Day 38) suggested that they were able to mount effective responses, resulting in inactivation of the virulent XDR *A. baumannii* although it was decreased to SBA KI=19.27 after injecting the lethal dose (1.5×10^8^ CFU/ml) which might be due to utilization of the previously formed antibody to kill the virulent *A. baumannii* injected by intraperitoneal route.

In a study by Shu *et al*. (2016) showed that, that the complement-mediated serum bacteriolysis effects in sera of mice from the first inoculation with inactivated whole-cell vaccine collected on day 7 and 12 showed only a slight incremental increase, however, the bacteriolysis effect in the sera collected on day 30 from immunized mice was much higher in comparison to the sera collected on the same day from control group mice.

These results suggest that inoculation of heat inactivated whole-cell vaccine stimulated an immune response that mediated significantly better clearance or bacteriolysis activity of XDR *A. baumannii*. Though, A single immunization did not significantly increase the complement mediated bacteriolysis activity, the results after the booster immunization of the vaccine group were able to mount effective responses, resulting in inactivation of the XDR *A. baumannii*. As a 4-fold or higher increase in antibody titre after vaccination is considered as a positive response, in comparison of the rising fold of IgG antibody titers of the EG group with PC group and analysis of SBA titres in post vaccinated sera showed that the IgG antibody titre were significantly increased after 2nd (7-fold) and 3rd inoculation (12-fold). On the other hand, 7-fold rise in SBA KI from 50% SBA titre was only shown after 3rd inoculation. These variations might be due to several factors including source and quantity of exogenous complements, bacterial strain, test sera, and antigen expression in targeted bacteria.

Although there was variation in the rising of titres, both studies showed effective killing of XDR *A. baumannii in vivo* and *in vitro* assay. These findings here are consistent with earlier studies, which suggest that a specific antibody response stimulated by a HI-WCV can aid in the clearance of *A. baumannii* and accord protection against *A. baumannii* infections (20). This vaccine development approach may potentially further the development of viable vaccines against XDR *A. baumannii* infections.

## Materials and Methods

### Sample collection

Clinical samples like endotracheal aspirate, blood, wound swab was collected from patients admitted in different departments, especially intensive care unit and burn unit of Dhaka Medical College Hospital.

### Phenotypic identification

*Acinetobacter spp*. was identified by observing colony morphology on blood agar (non-pigmented, white or cream-colored, smooth to mucoid colonies, 1-2 mm in diameter, non-hemolytic), on MacConkey agar (generally form colorless colonies), Gram staining (gram negative coccobacilli) and biochemical tests-oxidase test (negative), and catalase tests (positive), urease production (variable), indole test (negative), motility (non-motile) and citrate utilization test (positive) (21)

### Genotypic identification

PCR with *blaOXA-51-like* gene was used to identify *A. baumannii* (22) from the isolated *Acinetobacter spp*.

### Antimicrobial susceptibility test

Susceptibility was determined by modified Kirby-Bauer disc diffusion technique using Mueller-Hinton agar plates and zones of inhibition were interpreted according to Clinical Laboratory Standard Institute guideline (23). Criteria of the United States Food and Drug Administration (24) was used for the interpretation of zone of inhibition of tigecycline. The antimicrobial discs (Oxoid Ltd, UK) Amoxiclav (amoxicillin 20μg and clavulanic acid 10μg), piperacillin-tazobactam (100μg)/10μg), ceftazidime (30μg), ceftriaxone (30μg), cefepime (30μg), cefoxitin (30μg), aztreonam (30μg), ciprofloxacin (5μg), imipenem (10μg), amikacin (30μg), gentamycin (10μg), doxycycline (30μg), tigecycline (15μg) and colistin (8mg/ml) and *Escherichia coli* ATCC 25922 and *Pseudomonas aeruginosa* ATCC 27853 were used as control strains (25). The isolate that was resistant to all the 14 antibiotics was further isolated for preparation of HI-WCV.

### Preparation of HI-WCV

A loop of six XDR *A. baumannii* was inoculated into TSB and cultured overnight at 37°C and then centrifuged at 3,000g for one minute. The supernatants were discarded, and the cell pellets were washed twice with PBS. Subsequently, the bacterial concentrations were determined by serial dilution with normal saline and plating onto MacConkey agar plates followed by incubation at 37°C for 24 hours. The bacterial cells corresponding to 1×10^7^ CFU/ml were heat-inactivated in a water bath by incubating for 30 minutes at 56°C. Complete inactivation of the bacteria was confirmed by plating the supernatant onto Mueller-Hinton agar plates (19) and then stored at −20°C until inoculation.

### Mice and inoculation schedule

Eleven 6-8 weeks old male swiss albino mice were collected and kept under specific pathogen-free conditions with non-medicated diet. The mice were randomly divided into 3 groups where EG and PC group contained 4 mice and negative control (NC) group contained 3 mice.

### Inoculation schedule of mice

Three IM inoculations were performed on day 0, 14, 28 **(Figure 1A)** in the alternate thigh of the EG group of mice with 20μl of heat-inactivated whole-cell *A. baumannii* emulsified with 20μl CFA (1:1), and with 20μl of PBS emulsified with 20μl CFA (1:1) for the PC group of mice on the same day. The IM inoculation was done with an insulin syringe BD Ultra-Fine TM (31G). CFA (Sigma-Aldrich, Missouri, 50 United States) was used for the first inoculation. The prepared mixtures were inoculated after intraperitoneal injection of ketamine (ketamine 100 mg/kg) and chloroform by inhalation.

### Serum collection

Tail blood was collected after sterilization with 70% alcohol before inoculation and on 10^th^ day after each inoculation and about one ml blood was collected in a sterile test-tube by cardiac puncture from euthanized NC group, immediately after death from PC group and 14 days after the lethal challenge from EG group **(Figure 1A).**

### Detection of IgG antibody

Bacterial cells used in preparation of HI-WCV of *A. baumannii* (1×10^7^ CFU/ml), was used as antigen after sonication. Protein concentration was detected by nanodrop spectrophotometer (ThermoFisher Scientific). Immunoplates (Maxisorp, Nun) were coated with 100 μl/well of 10μg/ml antigen overnight at 4°C in the carbonate buffer. Subsequently, wells were blocked with 200μl blocking buffer (5% w/v non-fat dry milk in PBS-0.05% tween 20) and incubated at 37°C for 1-2 hour(s) after covering with adhesive plastic and washed 3 times. 100μl of diluted serum sample 1:100 in 1% bovine serum albumin in 1×PBS containing 0.1% Tween (PBST) were added to duplicate wells and incubated at 37°C for one hour. To each well, 100μl of a 1:5000 dilution of goat anti-mouse IgG peroxidase conjugated antibodies (Thermo Fisher Scientific, USA) was added and incubated at 37°C for one hour. The plates were washed three times with a wash buffer (1×PBST) post all incubations. The plate was developed using TMB, following 1M sulfuric acid addition to stop the reaction, and read at 450/630nm by ELISA plate reader (Biotek Inc., USA). Cut off value of OD was calculated by following formula:

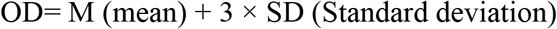

### Serum Bactericidal Antibody Assay

An XDR *A. baumannii* isolated stock was made by using PBS to determine the dilution (1:1,250) necessary to yield ~100 CFU/spot on Mueller-Hinton agar plates. The serum samples from EG of mice were heat-inactivated in a water bath by incubating at 56°C for 30 minutes. 3-4 weeks old two guinea pigs’ sera were used as a source of complement and to heat-inactive, serum was incubated at 56°C for 30 min. SBA was performed with some modifications (26). After diluting of the bacteria in 20ml of PBS according to the pre-determined optimal dilution factor (1:1250), 10μl of diluted bacteria was added to each well of the assay plate. 50μl of 20% solution heat-inactivated guinea pig complement was added to all wells in column 1 (control A wells) and 50μl of 20% solution native guinea pig complement was added to all wells in columns 2 through 12 (Control B and test sample wells). Plate was incubated at 37°C for 3 hours after brief mixing. 10μl of the reaction mixture was taken from each well onto a Mueller-Hinton agar plate, tilted and allowed the spots to run for ~1cm and incubated in the incubator upside-down for 16-18 hours.

Photograph of the plates was taken using a digital camera and transferred images to a computer to count the number of colonies in each spot. Calculation of Non - specific killing and 50% Killing Index was done by the following formula:

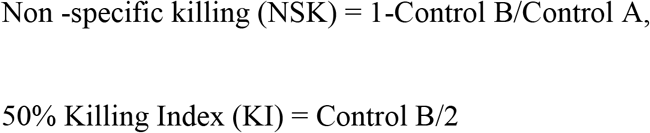

Because a serum dilution would rarely yield exactly this 50% KI value, it was interpolated from two sequential serum dilutions, one that kills less than 50% and one that kills more than 50%, **(Figure 2. a, b, c, d)**

The formula for calculating the interpolated SBA KI is shown below:

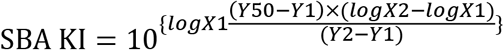

### Statistical analysis

The data from the antibody response, complement-mediated serum bacteriolysis were expressed as the mean± standard deviation from repeated assays. The data were subjected to two-way analysis of variance (ANOVA). Statistical significance was considered as *P*<0.05. All the statistical analyses were performed using Stata software version 14. (StataCorp LLC, USA).

## Ethical clearance

The animal experiments were performed in accordance with the protocols and guidelines approved by Dhaka Medical College Ethical Review Committee (MEU-DMC/ECC/2019/256). Informed written consent was obtained from all patients or legal guardians following a detailed explanation of the study.

## Acknowledgment

We thank Prof. Dr. S M Shamsuzzaman, Head of the department of Microbiology and Chairman of Ethical Review Committee of Dhaka Medical College for guiding us for vaccine evaluation and study as well as providing all laboratory support.

Dr. Nazmun Sharmin and Dr. Mahbub E Khoda performed vaccine design, laboratory work planning, sample processing and the animal experimentation. Dr. Mohammad Nazim Uddin performed data analysis and drafted the paper. All authors substantively revised and reviewed the paper and agreed to its contents.

